# AMPK/ULK1 activation downregulates TXNIP, Rab5, and Rab7 and inhibits endocytosis-mediated entry of human pathogenic viruses

**DOI:** 10.1101/2024.08.06.606943

**Authors:** Viktoria Diesendorf, Veronica La Rocca, Michelle Teutsch, Haisam Alattar, Helena Obernolte, Kornelia Kenst, Jens Seibel, Philipp Wörsdörfer, Katherina Sewald, Maria Steinke, Sibylle Schneider-Schaulies, Manfred B. Lutz, Mathias Munschauer, Jochen Bodem

## Abstract

Cellular metabolism must adapt rapidly to environmental alterations and adjust nutrient uptake. Low glucose availability activates the AMP-dependent kinase (AMPK) pathway. We demonstrate that activation of AMPK or the downstream Unc-51-like autophagy-activating kinase (ULK1) inhibits receptor-mediated endocytosis. Beyond limiting dextran-uptake, this activation prevents endocytic uptake of human pathogenic enveloped and non-enveloped, positive and negative-stranded RNA viruses, such as yellow fever, dengue, tick-borne encephalitis, chikungunya, polio, rubella, rabies lyssavirus and SARS-CoV-2 not only in mammalian and insect cells but in precision-cut lung slices and neuronal organoids. However, receptor presentation at the cytoplasmic membrane was unaffected, indicating that receptor-binding remained unaltered and later steps of endocytosis were targeted. Indeed, AMPK pathway activation reduced early endocytic factors TXNIP, Rab5 and the late endosomal marker Rab7 amounts. Furthermore, AMPK activation impaired SARS-CoV-2 late-replication steps by reducing viral RNAs and proteins and the endo-lysosomal markers LAMP1 and GRP78, suggesting a reduction of early and late endosomes and lysosomes. Inhibition of the PI3K and mTORC2 pathways, which sense amino acids and growth factor availability, promotes AMPK activity and blocks viral entry. Our results indicate that AMPK and ULK1 emerge as restriction factors of cellular endocytosis, impeding the receptor-mediated endocytic entry of enveloped and non-enveloped RNA viruses.

## Introduction

Cellular metabolism must be tightly and rapidly adjusted to the availability of nutrients and growth factors. The cell adapts to environmental alterations by activating and inhibiting kinase signalling pathways. Elevated levels of amino acids and growth factors trigger the activation of the phosphatidylinositol 3-kinase (PI3K)/Akt pathway, positively regulating transcription by the RNA polymerases I and II, as well as translation of 5’ terminal oligopyrimidine tract (TOP) mRNAs. Additionally, the mammalian target of rapamycin (mTOR), a downstream kinase, regulates lysosomal and endoplasmic reticulum function while inhibiting degenerative processes like lysosome acidification and autophagy.

Conversely, a glucose undersupply, resulting in reduced cellular ATP levels, is sensed by the liver serine-threonine kinase B1 (LKB1) activating the AMP-activated protein kinase (AMPK). The AMPK is a heterotrimeric complex that acts as the core energy sensor of the cells (reviewed in ^1^). AMPK phosphorylates the Unc-51-like autophagy-activating kinases (ULK1), promoting autophagy (Fig. 1A). Simultaneously, the mTORC1 complex is inactivated by Raptor phosphorylation, resulting in lysosome acidification due to the relief of the suppression of V-ATPase by binding to the mTORC1 complex.

**Figure 1.**
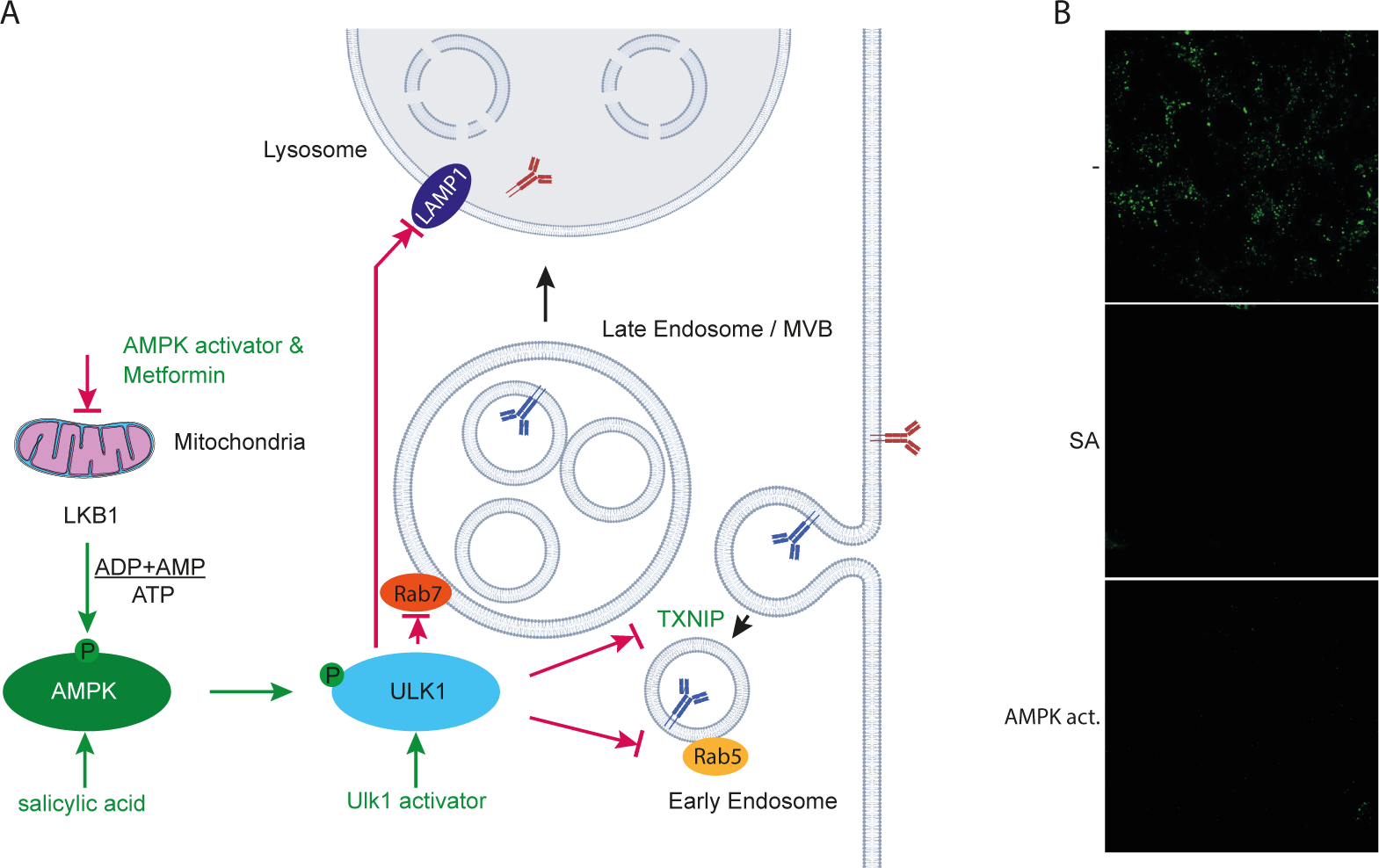
AMPK activation inhibits dextran endocytosis. **(A)** Scheme of AMPK kinase pathways and mechanism of endosome maturation. green: activators; red: inhibitors; green arrows: activation; red arrows: inhibition. **(B)** Confocal microscopy of cells incubated with 3 mM SA or 100 µM AMPK activator and 2.5 mg/ml fluorescein isothiocyanate-dextran. The cells were fixed after 2 h.

As suggested by prior work, the AMPK pathway may regulate endocytosis-dependent glucose uptake ^1^. In addition, aspirin (acetylsalicylic acid (ASA)) was shown to activate AMPK ^2, 3^ and, in turn, inhibit the vitamin C uptake in mice ^4^. Furthermore, active AMPK negatively regulates glucose transporter 1 (GLUT1) endocytosis in low ATP levels to ensure glucose resupply. Under physiological conditions and in response to high glucose availability, thioredoxin-interacting protein (TXNIP)^5^, a member of the α-arrestin family, hinders glucose uptake by binding to GLUT1 and promoting its internalisation, ubiquitinoylation, and delivery to the lysosome ^6, 7^. AMPK phosphorylates TXNIP at the serine residue 308, leading to its rapid degradation, thus enhancing glucose uptake by inhibiting the recycling of GLUT1 (Fig. 1A). Similarly, Akt phosphorylates TXNIP at the same serine residue to ensure glucose uptake under conditions of high amino acid and growth factor supply^8^.

TXNIP is also regulated by miR-183, which downregulates its expression. High nutrient availability activates the p65 subunit of the NF-kappa B (NF-κB) transcription factor through the Akt/mTORC1/2 kinase pathway, upregulating TXNIP via enhanced histone deacetylase 2 (HDAC2) expression ^9, 10^. Thus, the Akt/mTORC1/2 kinase pathway alleviates the TXNIP-mediated suppression of GLUT1 endocytosis through Akt and NF-κB p65 ^10^.

Organelles in the endocytic system are defined by their Rab GTPases, which serve as building platforms for the specific factors of the organelles^11^. Rab5 is associated with the outer membrane early endosome (EE) and is involved in endosome fusion (Fig. 1A) ^12^. Rab4 and 11 control the endosome recycling to the plasma membrane. Chemical AMPK activation enhances Rab4 promoter activity ^13^. Rab5 is activated by its guanine nucleotide exchange factor Rabex5, which binds to the ubiquitinylated cargo. Rab5 recruits the antigen 1 (EEA1), a tethering protein required for the fusion of endocytic vesicles with EEs, and the hexameric CORVET tethering complex, functioning in the fusion of endocytic vesicles and among EEs ^14,15^. CORVET binds to Rab5 and participates in the fusion of endocytic vesicles and EEs. During the switch from Rab5 to Rab7, Rab5 and Rabex5 are released from the endosome, and the Rab5-specific GTP-hydrolysis-activating protein inactivates the remaining Rab5 ^16^. The expression of constitutively inactive Rab7 resulted in cargo enrichment in the EEs ^17^, probably by blocking the endosomal pathway. However, Rab7 is essential for the transport from late endosomes (LE) to lysosomes (Fig. 1A), where Rab7 and its downstream effector Rab-interacting lysosomal protein recruit the V-ATPase and control the stability of the V1G1-vATPase subunit ^18^. Eventually, the LE fuses with the lysosome, releasing its content into the lysosomal lumen (Fig. 1A).

Recently, it has been reported that the Rift Valley fever virus (RVFV), an animal pathogen, is inhibited by AMPK activation. It was concluded that an early step in the viral replication cycle is inhibited by AMPK, suggesting that RVFV infection is restricted due to inhibition of fatty acid biosynthesis ^19^. Previously, it has been shown that ASA and its metabolite salicylic acid (SA) inhibit SARS-CoV-2, influenza A, and rhinovirus replication in cell culture cells and human precision-cut lung slices (PCLS) ^20–22^. In a more extensive retrospective cohort study, SARS-CoV-2 patients receiving ASA required less oxygenation on admission than patients not receiving ASA ^23^. Here, we investigate how signalling cascades related to nutrient availability and potential AMPK activation by ASA and SA impact the replication of flaviviruses, including yellow fever (YFV), dengue (DENV), tick-borne encephalitis virus (TBEV), the alphavirus chikungunya, the coronavirus SARS-CoV-2, and rabies lyssavirus. Endemic areas of these viruses continue to expand. DENV is now endemic in over 100 countries across WHO regions, with the number of cases increasing from 505,430 in 2000 to 5.2 million in 2019 globally and 2.8 million cases reported in the Americas alone in 2022. While an attenuated vaccine for DENV has been approved, YFV still causes 200,000 diagnosed cases and 30,000 deaths annually despite the vaccine’s introduction in 1937. Similarly, despite approved rabies vaccines and postexposure prophylaxis, between 50,000 to 70,000 fatal canine rabies infections occur every year ^24^. These findings indicate that developing antiviral therapies is still an essential health issue.

It has been observed that SA efficiently activates AMPK by direct interaction ^3^. While our previous work demonstrated the inhibitory effect of ASA/SA on SARS-CoV-2, the underlying mechanism remained unexplored. If the inhibition of SARS-CoV-2 by ASA/SA is indeed mediated through AMPK activation, it follows that direct activation of AMPK might also inhibit other viruses. In the present work, we investigated whether an AMPK activator could similarly suppress the replication of different viruses despite the differences in their replication cycle. We provide evidence that AMPK activation efficiently blocks the replication of multiple human pathogenic RNA viruses.

## Results and Discussion

### AMPK activation suppresses receptor-mediated dextran-uptake

Since it has been shown that GLUT1/4 endocytosis and RVFV entry are, although unrelated, inhibited by AMPK activation, we hypothesised that AMPK activation impedes, more broadly, receptor-mediated endocytosis. Thus, we sought to investigate whether AMPK activation inhibits the endocytosis of fluorescently labelled dextran, a marker of receptor-mediated endocytosis. After dextran was taken up, the Huh-7 cells were incubated with SA or an AMPK activator, and dextran was added to the medium for 2 h. The cells were fixed and subsequently analysed by confocal microscopy (Fig. 1B). Indeed, activation of AMPK by both compounds suppressed the dextran uptake, indicating that the suppression of receptor-mediated endocytosis was impaired.

### Activation of the AMPK inhibits flaviviruses at viral entry

Since other (+)-stranded RNA viruses commonly use endosome-derived replicative compartments derived from the endosomes, the endoplasmic reticulum, and lysosomes as well, we analysed whether AMPK activation by ASA/SA inhibits DENV, TBEV, and YFV replication. Huh-7 (DENV, YFV) or human glioma cells NCE U373 (TBEV) cells were pretreated with 3 mM ASA/SA, 100 µM AMPK or ULK1 activators for 2 h prior to infection (Fig. 2A). The AMPK activator does not directly interact with AMPK but inhibits the mitochondrial complex I, reducing cellular ATP levels and activating AMPK ^25^. The viruses were removed 2 h after infection by a medium exchange. Cell culture supernatants were collected after 2 (YFV, TBEV) or 3 (DENV) days; viral RNAs were isolated and quantified by RTqPCR (Fig. 2C to G). All infection experiments in this manuscript were performed three times in triplicate. Accumulation of both TBEV and DENV RNAs was inhibited by more than 2 orders of magnitude (log_10_), showing that the ASA/SA treatment suppressed these viruses efficiently. Treatment with the AMPK activator inhibited TBEV replication by approximately 1.3 log_10_ and YFV by more than 3 log_10_, indicating that the AMPK activation was sufficient to repress viral replication (Fig. 2D and F). The activation of the downstream ULK1 suppressed YFV replication even further, showing that inhibition of viral replication was mediated through the ULK1.

**Figure 2.**
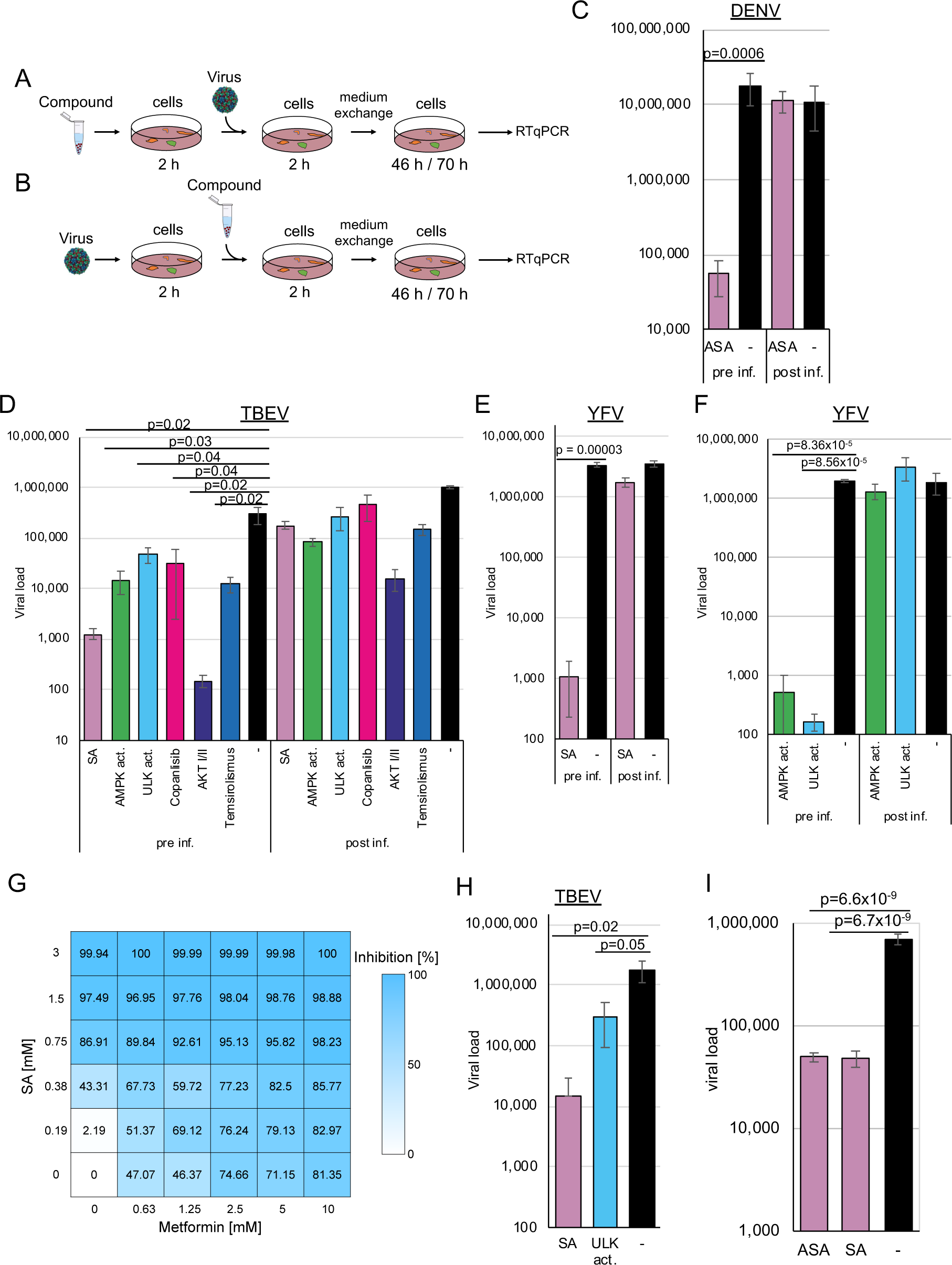
AMPK and ULK1 activation or PI3 kinase inhibitors inhibit flavivirus entry. **(A)** Cells were pre-treated with the AMPK, ULK1 activators or kinase inhibitors for 2 h and infected with (pre inf.) **(C)** DENV, **(D)** TBEV or **(E** and **F)** YFV, or **(B)** infected for 2 h and treated (post inf.). **(G)** Inhibition of YFV replication by combinations of SA and metformin. **(H)** SA and ULK1 activators suppress TBEV replication in neuronal organoids. **(I)** AMPK activation blocks YFV replication in SF9 insect cells. SA inhibits YFV in insect cells. Viral replication was determined by RTqPCR.

Metformin, a commonly used antidiabetic drug, was also shown to activate AMPK ^26, 27^. Given that high doses of SA are necessary for AMPK activation, we aimed to investigate whether a similar activation of AMPK could be achieved through the combination of metformin and SA. We anticipated that both compounds would exhibit additive effects. Cells were treated with increasing concentrations of SA from 0.185 to 3 mM and metformin from 0.625 to 10 mM and infected with YFV (Fig 2G). The EC_50_ for SA was calculated to be 0.42 mM, and for metformin with 1.91 mM. Concentrations of 1.5 mM SA suppressed YFV replication by 97.49%, while 10 mM metformin inhibited viral replication by 81.4%. The analyses of drug synergism with the SynergyFinder software version 3.0 ^28^ showed that the compounds act additively, leading to an inhibition of more than 98 % with 2.5 mM metformin and 1.5 mM SA or 10 mM metformin and 0.75 mM SA. Consequently, the combination of metformin and SA either results in enhanced inhibition of viral replication or requires lower compound concentrations, thereby achieving comparable suppression, suggesting that the decreased SA concentrations would lead to adverse effects on fever in patients.

Recently, we and others have observed that even direct antivirals can act in a cell-type-specific way ^29–31^. Consequently, we aimed to validate our findings regarding TBEV using patient-relevant systems, specifically employing human stem cell-derived 3D neuronal organoid models. The organoids were exposed to 3 mM SA or 100 mM ULK1 activator and subsequently infected with TBEV. Cell culture supernatants were collected 2 days after infection, and viral RNAs were quantified by RTqPCR (Fig. 2H). SA downregulated TBEV replication by more than 2 log_10_ and the ULK1 inhibitor by 6-fold. These results indicate that activation of AMPK inhibits TBEV replication even in this patient-near model system. Since mosquitoes transmit YFV, we investigated whether the AMPK activation influences viral replication in insect SF9 cells. We observed the antiviral effects of ASA and SA in these cells, providing evidence that the mechanism is conserved (Fig. 2I).

Next, we aimed to determine which replication step was targeted by AMPK activation. To discern whether the compounds inhibited viral replication at an entry of the post-entry step, drugs were added either 2 h (Fig. 2A) before (pre-treatment) or 2 h after viral infection (Fig. 2B). For the latter setting, 3 mM ASA or SA, 100 µM AMPK activator (Fig. 2D to E), or 100 µM of the ULK1 were used (Fig. 2F). The cells were incubated for 46 (YFV, TBEV) or 70 h (DENV). Viral RNAs were isolated and quantified by RTqPCR (Fig. 2). We expected a lack of inhibition with AMPK activators similar to the untreated controls if AMPK activation affected only viral entry since the compounds were added after infection. We observed inhibition of viral replication by more than 2 log_10_ when compounds were added 2 h before viral entry. However, treatment with the compounds after infection did not reduce DENV infection, indicating that compound-mediated AMPK activation interfered with DENV entry. Similar experiments with YFV and TBEV yielded comparable results, suggesting that AMPK activation inhibits viral entry similarly. Remarkably, the pharmacologic inhibition of receptor-mediated viral entry by the ULK1 activator suggested that this downstream kinase also regulates flavivirus entry, revealing a previously unrecognised role in the viral infection process.

### AMPK activation inhibits Enteroviruses, Alphavirus, and Rubella virus entry

Having observed that flavivirus entry is sensitive to AMPK activation, we also sought to analyse whether this phenomenon applies to other plus-stranded RNA viruses such as rubella, chikungunya, polio- and enterovirus 71. Like RVFV, the chikungunya virus enters the cell via the Mxra8 receptor. The myelin oligodendrocyte glycoprotein is a rubella virus receptor, the scavenger receptor class B member 2 or P-selectin glycoprotein ligand-1 for enterovirus 71, while polioviruses use CD155 for receptor-mediated endocytosis ^32–35^. Thus, we expected that AMPK activation would inhibit early infection steps only if the entry is regulated by cellular and not by viral factors.

We compared viral replication in cells pretreated for 2 h with that in cells infected 2 h before treatment to further analyse if the activation of AMPK by SA suppresses viral entry of these viruses in cells. The cells were either preincubated for 2 h with 3 mM SA, infected with the viruses, or infected 2 h before the treatment. Viral replication was determined by RTqPCR 2 d after infection. Activation of AMPK suppressed chikungunya (SA: 2.14 log_10_, p=1.4×10^-5^; AMPK activator 0.9 log_10_, p=2.4×10^-5^) (Fig. 3A), rubella (2.61 log_10_, p=0.001) (Fig. 3B), EV71 (SA: 2.81 log_10_, p=0.002) (Fig. 3C) and poliovirus (2.11 log_10_, p=0.002) (Fig. 3D) when cells were preincubated with the compounds. AMPK activation did not significantly reduce viral replication, indicating that virus entry is inhibited, similar to our results with flaviviruses and dextran.

**Figure 3.**
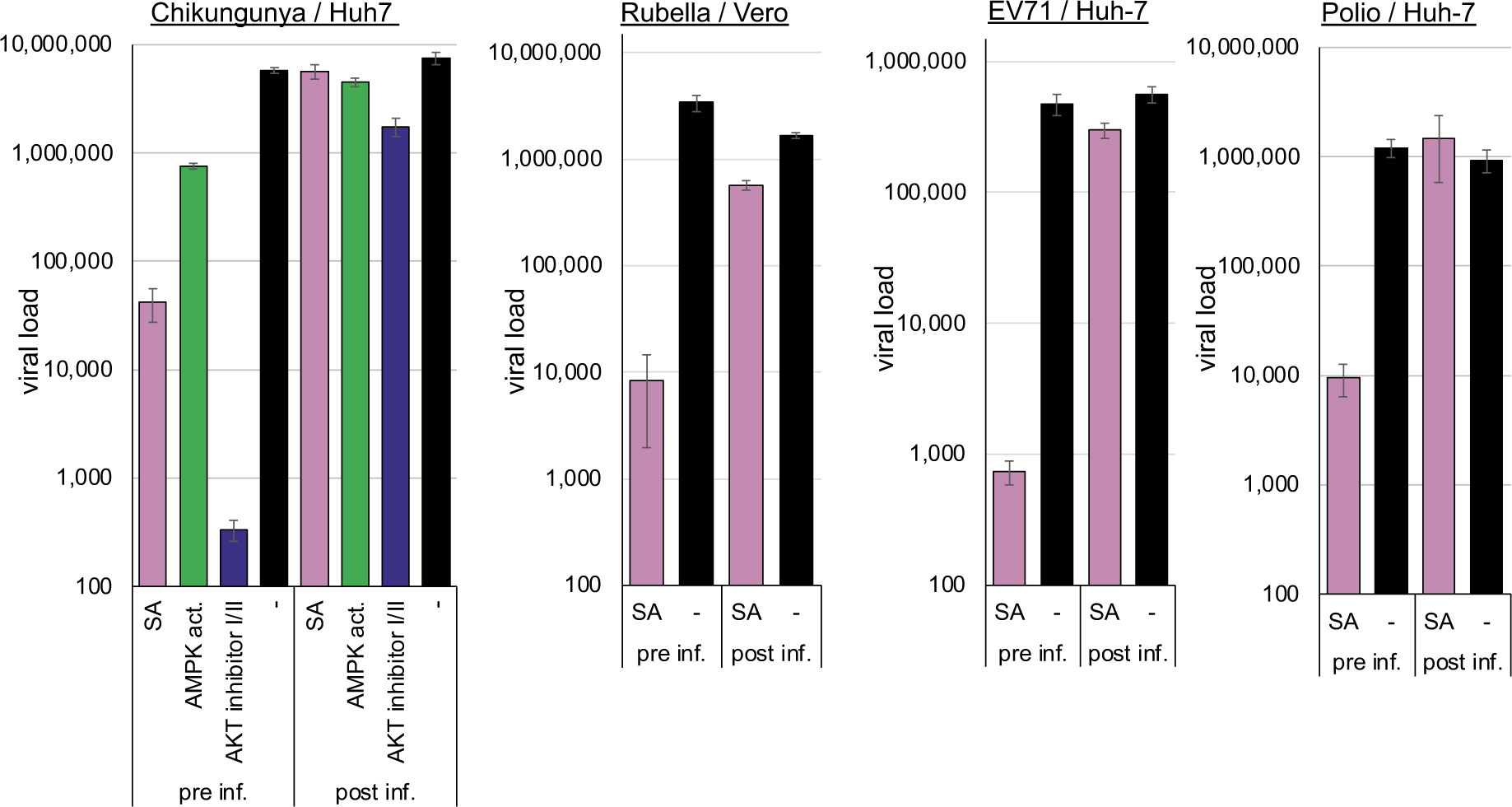
AMPK activation inhibits Chikungunya, Rubella, EV71, and Poliovirus entry. Cells were preincubated for 2 h with the compounds and infected with Chikungunya (pre inf.) or infected and supplemented with the compounds 2 h after infection (post inf.). Released viral genomes were quantified 48 h after infection. Cell lines and viruses are depicted above the panel.

### AMPK activation does not inhibit SARS-CoV-2 entry into Calu-3 but into Vero cells

Recently, we showed that ASA and SA inhibit SARS-CoV-2 replication in cell culture and human PCLS ^22^. Quantifying viral RNA in infected cells at different time points did not support a correlation between the inhibition of viral replication and the interference of viral entry in Calu-3 cell cultures. However, since SARS-CoV-2 can enter cells via direct fusion with the plasma membrane or endocytosis using the same receptor ^36, 37^, we reanalysed the influence of SA on entry with a modified experimental approach using Calu-3 or Vero cells (Fig. 4A and B). Calu-3 cells express high amounts of the TMPRSS2 protease and, hence, promote viral entry by fusion at the plasma membrane. Entry into Vero cells is preferentially endocytic, which we enforced by adding the TMPRSS2 inhibitor nafamostat (Fig. 4B). This experiment allowed the determination of AMPK-mediated influences on the entry pathways since the virus and the receptor are invariant.

**Figure 4.**
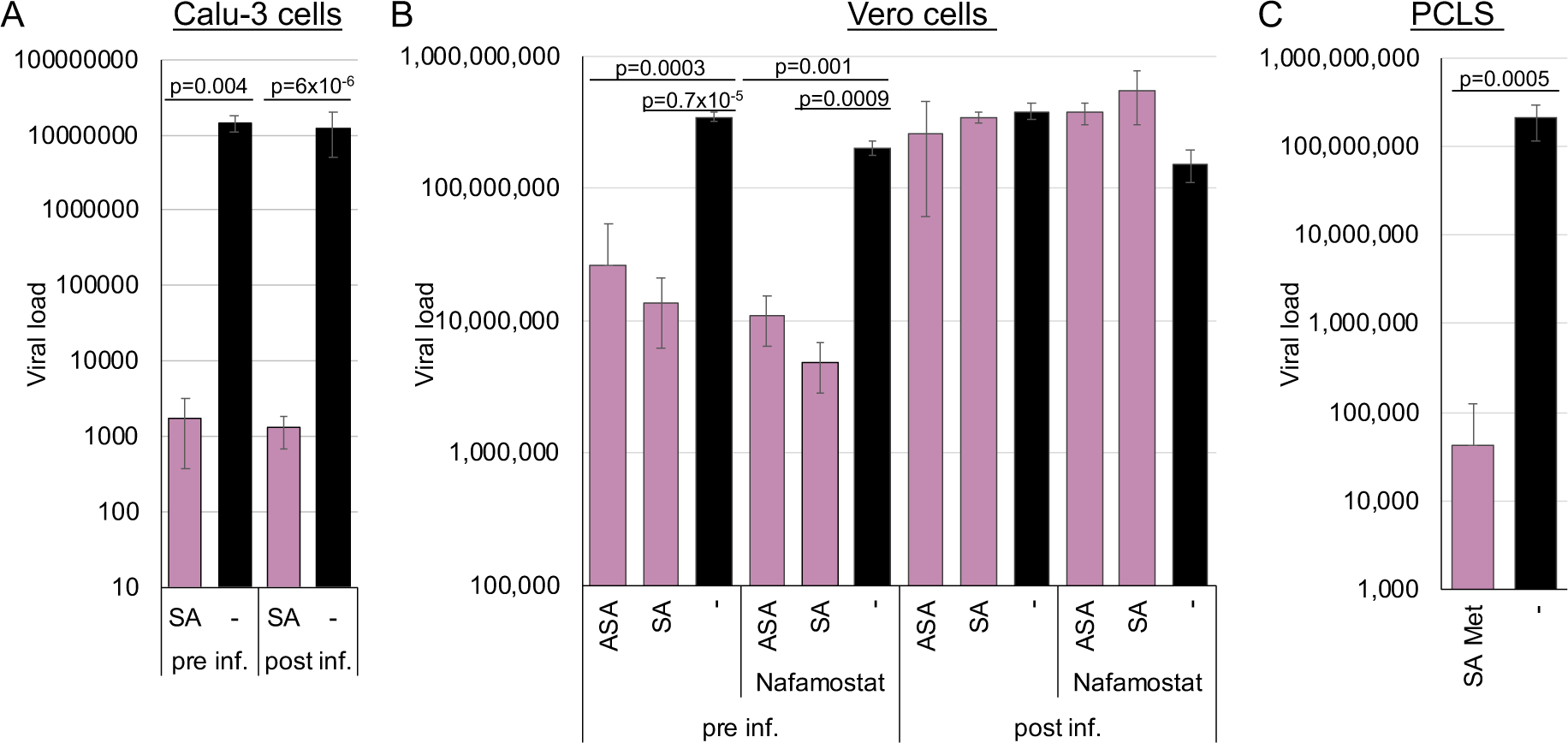
AMPK activation inhibits SARS-CoV-2 in Calu-3 and Huh-7 cells but does not influence viral entry in Calu-3 cells. **(A)** Calu-3 cells or **(B)** Vero cells were preincubated for 2 h with 3 mM SA and infected (pre inf.) or infected and supplemented with 3 mM SA 2 h after infection (post inf.). Nafamostat was added to ensure entry by endocytosis. **(C)** Treatment with 1.5 mM and 5 mM metformin inhibits SARS-CoV-2 in precision-cut lung slices. Released viral genomes were quantified 72 h after infection.

The Calu-3 cells were either preincubated for 2 h before infection with 3 mM SA or infected with SARS-CoV-2, and 3 mM SA was added 2 h after infection. In the first case, viral replication should be inhibited if SA targets viral entry, while in the second case, viral entry should not be affected. The medium was exchanged after 6 h, and viral genomes were quantified after 72 h by RTqPCR (Fig. 4). Virus replication in Calu-3 was independent pre- or post-incubation, underlining that AMPK did not target the entry via direct fusion at the plasma membrane and confirmed our previous findings.

To further analyse if AMPK activation by a combination of SA and metformin inhibits SARS-CoV-2 in a patient-near system, we treated human PCLS with 5 mM metformin and 1.5 mM SA (Fig. 4C). The combination effectively suppressed viral replication, showing that AMPK activation inhibits SARS-CoV-2 in a patient-near system.

A similar experiment with Vero cells showed that AMPK activation by ASA or SA before entry inhibited SARS-CoV-2 replication. However, the antiviral effect was completely abolished when viral entry occurred before AMPK activation, showing that ASA and SA specifically target endocytic viral entry, not receptor recognition or binding. Further inhibition of TMPRSS2 by nafamostat decreased viral replication slightly, indicating that the TMPRSS2 activity on Vero cells is low. These results suggest that AMPK activation can inhibit SARS-CoV-2 entry in cells expressing low amounts of TMPRRS2 and explain the observed correlation of fasting blood glucose with the severity of COVID-19 in patients even without diagnosed diabetes ^38, 39^. In addition, AMPK activation initiates a second antiviral mechanism blocking SARS-CoV-2 replication in Calu-3 cells.

### AMPK activation inhibits entry of negative-stranded RNA viruses, such as rabies virus, and supports rabies postexposure prophylaxis

Since we have shown that the endocytic entry of (+)-stranded RNA viruses is AMPK dependent, we investigated whether the negative-stranded RNA virus entry of the vesicular stomatitis virus and rabies virus is affected by AMPK activation. In particular, if ASA/SA inhibited rabies replication, they could be added to the postexposure prophylaxis with antisera to prevent rabies infections. Inhibition and entry analyses were conducted as described above. BHK-21 or mouse NA 42/13 cells were either pretreated with 3 mM ASA/SA, 100 µM AMPK, or ULK1 activators and infected with the rabies vaccine (BHK-21 cells, Fig. 5A) or *wild-type* (NA 42/13 cells, Fig. 5B) strain or infected and incubated with the compounds after 2 h. Supernatants were harvested after 75 h, and viral genome copies were determined by RTqPCR. Pretreatment with ASA or SA inhibited replication of the rabies vaccine strain more than 2 log_10_ (ASA: 2.2 log_10_, p=0.04; SA: 2.4 log_10_, p=0.04). AMPK activation inhibited entry of the wild-type rabies virus by approx. 1.5 log_10_ (ASA: 1.6 log_10_, p=0.002, AMPK activator: 1.5 log_10_, p=0.002) into NA 42/13 cells while the ULK1 activator reduced entry by 0.5 log_10_ (p=0.013). Thus, activation of AMPK downregulates rabies virus entry.

**Figure 5.**
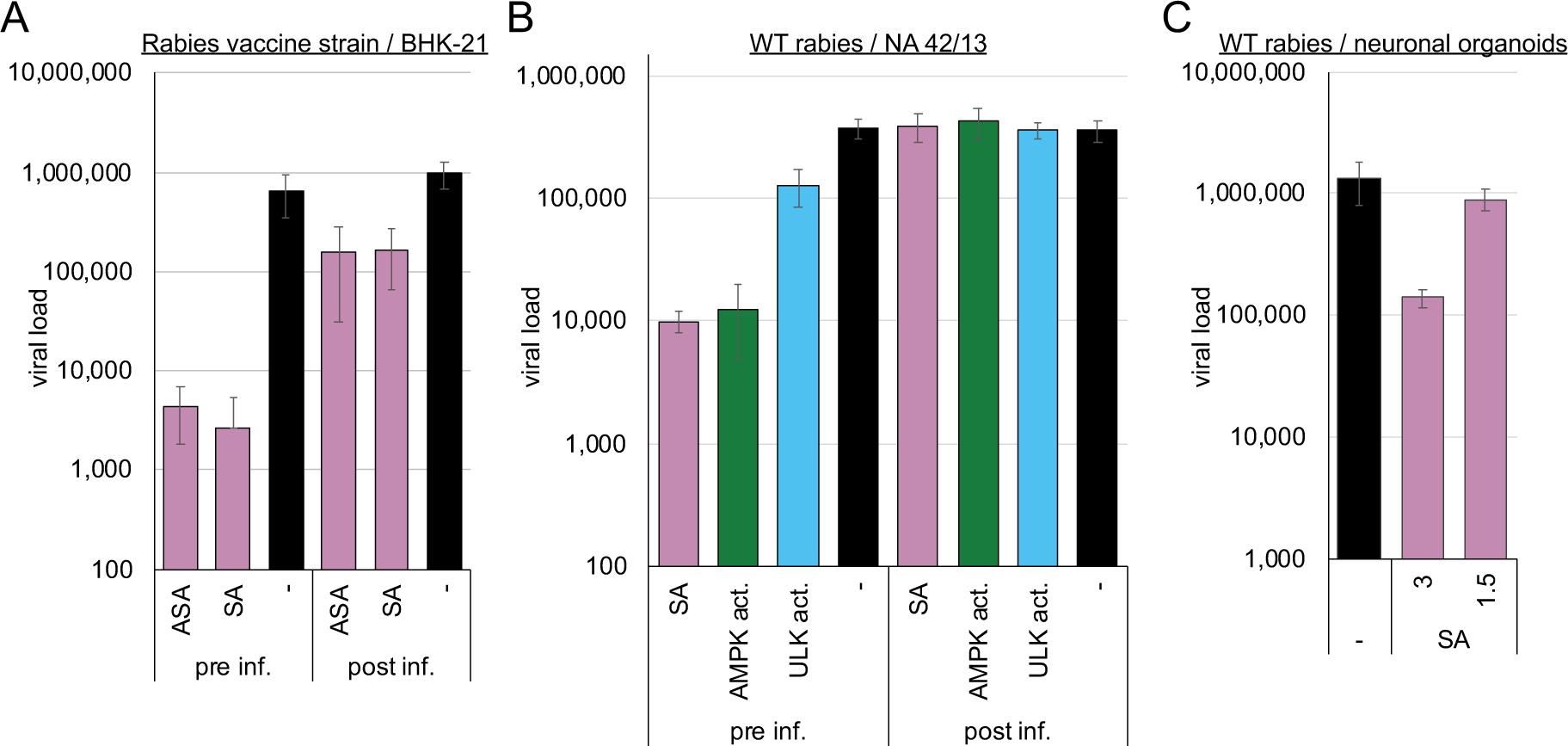
AMPK and ULK1 activation inhibits rabies vaccine and *wild-type* strains. **(A)** BHK21, **(B)** NA42/13 cells, or **(C)** neuronal organoids were preincubated for 2 h with the compounds (3 mM ASA/SA, 100 µM AMPK or UKL1 activator) and infected with **(A)** rabies vaccine or **(B & C)** *wild-type* strain (pre inf.) or infected and supplemented with compounds 2 h after infection (post inf.).

The AMPK-induced inhibition of viral replication was confirmed in human stem cell-derived 3D neuronal organoid models. The organoids were exposed to 1.5 or 3 mM SA and subsequently infected with wild-type rabies virus. The medium was exchanged after 24 hours to remove the nonincorporated virus. Supernatants were collected after 3 d of infection, and viral RNAs were quantified by RTqPCR (Fig. 5C). 3 mM SA reduced viral genome amounts by 1 log_10_ (p=0.05). Similar inhibition of the vesicular stomatitis virus was obtained after AMPK activation with SA (Fig. S1).

AMPK activation inhibited rabies *wild-type* and vaccine strain entry and replication. Thus, adding an AMPK activator to the approved antisera for postexposure prophylaxis or, if antisera are unavailable, their use to clean the bite wounds might be beneficial for exposed patients.

### AMPK activation does not influence receptor presentation but downregulates TXNIP, Rab5, and Rab7 and inhibits virus entry

We further investigated the mechanisms involved in the AMPK-mediated downregulation of viral entry. Viruses, such as DENV, TBEV, YFV, chikungunya, rabies, VSV, and SARS-CoV-2 or molecules, such as dextran, utilise specific entry receptors and should be regulated differently. Consequently, AMPK activation is hypothesised to impede viral entry and, more broadly, receptor-mediated endocytosis. First, we determined if AMPK or ULK1 activation changed the receptor presentation at the plasma membrane. We decided to analyse the poliovirus receptor CD155 and the VSV low-density lipoprotein receptor (LDL) entry receptor on MT4 cells by FACS ^40^ to address negative and positive-stranded viral receptor proteins. The LDL receptor supports the entry of Getah, Semliki Forest and Bebaru viruses besides VSV ^41^. MT4 cells were treated with the AMPK or ULK1 activator for 2 h and stained with the respective rabbit-anti-CD155 or mouse-anti-LDL receptor antibodies. FACS analyses showed that the amounts of both receptors were not changed by AMPK or ULK1 activation (Fig. 6A and S2), indicating that changes in the receptor concentration are not responsible for the observed entry inhibition.

**Figure 6.**
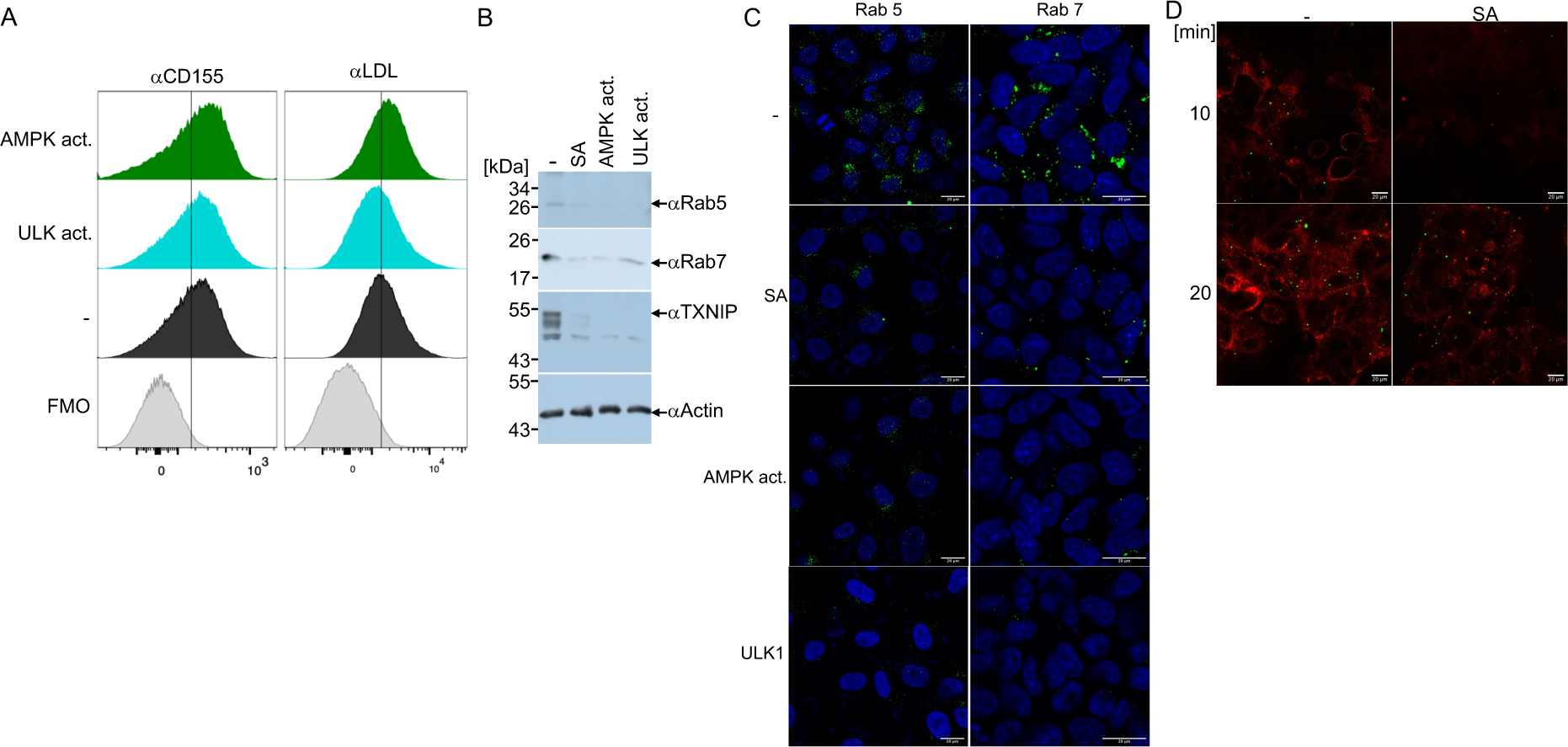
AMPK activation does not influence the surface expression of CD155 or the LDL receptor, but it reduces cellular TXNIP, Rab5, and Rab7 and inhibits viral endocytosis. **(A)** FACS analyses of the CD155 and LDLR expression at the plasma membrane. FMO: control without the 1^st^ antibodies. **(B)** Western blotting analysis of TXNIP, Rab5 and Rab7. **(C)** Confocal microscopy of Rab5 and Rab7. **(D)** Confocal microscopy (viruses, green; cellular membranes, red) of SA treatment and YFV infection. Pre-treated Huh-7 cells were infected with fluorescently labelled YFV for 10- or 20-min. Scale bar: 20 µm.

Generally, it is known that the α-arrestin TXNIP is involved in regulating the endocytosis of GLUT1. TXNIP is negatively regulated by AMPK through phosphorylation, promoting its degradation and causing inhibition of GLUT1 endocytosis. To analyse whether AMPK-mediated inhibition of viral entry is related to TXNIP degradation. Huh-7 cells were incubated with SA for 24 h or an AMPK activator for 2 h, and TXNIP expression was visualised by Western blotting using specific antibodies (Fig. 6B). The activation of AMPK decreased TXNIP amounts, suggesting that the first endocytosis steps are already inhibited, similar to the inhibition of GLUT1.

The Ras-related protein Rab5 marks the EE. It is enriched in the EEs and is later the primary regulator of the conversion to the LEs, where the Rab5-GTP recruits Rab7 to the endosome. Thus, EEs are marked by Rab5, while during the transition from the EEs to the LEs, Rab5 is replaced by Rab7. Analyses of the Rab5 and 7 amounts by Western blotting revealed that both proteins are downregulated by AMPK and, surprisingly, by ULK1 activation (Fig. 6B), indicating that ULK1 influences the Rab5-positive EEs. These results were confirmed by confocal microscopy with Rab5 and Rab7-specific antibodies (Fig. 6C). The Vero cells were treated for 2 h with the AMPK or ULK1 activators, fixed and stained. Confocal microscopy revealed that AMPK activation by SA or AMPK activator reduced Rab5 and Rab7 expression. Moreover, ULK1 activation also decreased the number of LEs and EEs. Thus, the AMPK pathway regulates EEs and LEs in addition to the Rab4-mediated endosome recycling that has been previously reported ^13^. Considering our results, the upregulation of Rab4 might compensate for the downregulation of Rab5 in EEs and Rab7 in LEs.

Next, we sought to show the effects of AMPK activation using fluorescently labelled YFV. Cells were infected with concentrated YFV at high MOI (MOI∼10). After 1 day, the medium was exchanged for medium containing ATTO 643 DOPE, a compound staining cellular membranes. Cell culture supernatants were harvested after 12 h, and viruses were pelleted by ultracentrifugation, resulting in a coloured pellet. Huh-7 cells were treated with SA for 2 h; then, the medium was replaced with a medium containing a cytoplasmic membrane cell stain (BioTracker 400 blue, Sigma-Aldrich) but no compounds. This dye is incorporated by endocytosis. The cells were infected with the coloured viruses for 10 or 20 min and fixed with paraformaldehyde. Confocal microscopy revealed that YFV could not be detected in cells treated for 10 min with SA, indicating that the viruses were not endocytosed (Fig. 6D). The membrane dye was not incorporated, suggesting a general block of endocytosis. However, after 20 min of infection without the compound, virus binding was restored, proving the inhibition was reversible. The results indicate that the AMPK activation blocks already the first step of endocytosis, as expected by the decreased amounts of TXNIP. Similarly, cellular Rab5 and 7 were reduced, indicating a reduction in the later steps of endocytosis.

### AMPK activation modifies the endo-lysosomal SARS-CoV-2 replication compartment

Since Rab7 recruits the V-ATPase required for the conversation of endosomes to the lysosome, we analysed whether AMPK activation inhibited SARS-CoV-2 replication at post-entry steps. First, we determined the influence of AMPK activation on viral gene expression in these cells. Calu-3 cells were incubated with ASA and infected with SARS-CoV-2. Cells were lysed 3 days after infection, and viral N- and spike protein amounts were detected by Western blotting analysis. In addition, cell culture supernatants were collected, and viruses were concentrated by ultracentrifugation through a sucrose cushion. The treatment with AMPK activation reduced the non-processed S protein below the detection level (Fig. 7A), while S and S2 were absent in the supernatant. However, detectable amounts of the N proteins were found in the concentrated viruses.

**Figure 7.**
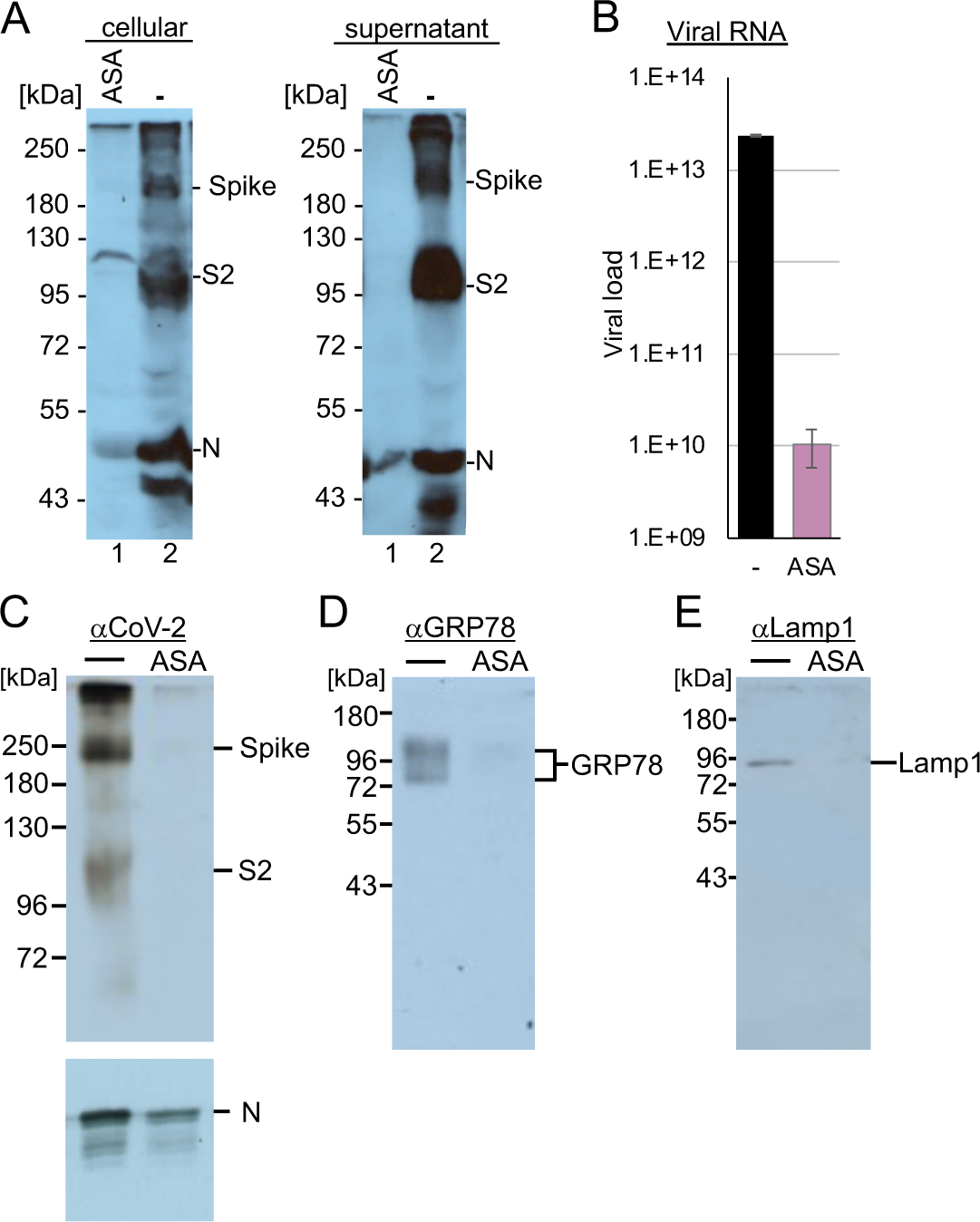
AMPK activation reduces (A) spike, nucleocapsid protein (N) expression and lysosomal amounts of (B) viral RNA, (C) spike protein, (D) endoplasmic marker GRP78 and the lysosomal marker protein Lamp1. **(A)** Calu-3 cells were incubated with ASA and infected with SARS-CoV-2 for 72 h. Infected cells and viral supernatants were analysed with anti-SARS-CoV-2 S2 and anti-SARS-CoV-2 N protein antibodies. **(B)** RTqPCR quantification and **(C to E)** Western blotting analyses of the lysosomal fraction of SARS-CoV-2 infected Calu-3 cells.

SARS-CoV-2 replication extensively uses the lysosomal compartment, and AMPK also controls lysosomal function and the endosomal-lysosomal compartment. Recently, we showed that the serotonin reuptake inhibitor fluoxetine traps SARS-CoV-2 in the lysosomes ^42^. Here, we sought to analyse whether AMPK activation modifies the replication compartment similarly. Calu-3 cells were incubated with ASA and infected with SARS-CoV-2 at an MOI of 10. After 24 h, the cells were lysed, and the lysosomes were isolated. The lysosomal pellets were dissolved in PBS, and viral RNAs and marker proteins were analysed. We determined a reduction of more than 3 log_10_ of viral RNAs and S protein levels almost undetectable in the cells treated with ASA compared to that in the untreated control (Fig. 7B and C), indicating that AMPK activation inhibited viral transcription and genome replication, resulting in less S and N proteins. Analyses of the endoplasmic reticulum marker GRP78 revealed that the expression of this protein, similar to that of the lysosomal marker protein LAMP1, was reduced in our lysosomal fraction (Fig. 7D and E). These results indicate that AMPK activation with ASA leads to fewer replicative lysosomal compartments, similar to the observed reductions of Rab5 and 7 amounts. Our results suggest that the ASA treatment modifies endosomal, lysosomal, and SARS-CoV-2 replication compartments.

### The PI3K/Akt kinase pathway is essential for flavivirus and SARS-CoV-2 replication

PI3K/Akt pathway promotes viral endocytosis by activating mTORC1 and silencing AMPK/ULK1. The AMPK activation antagonises the PI3K/Akt pathway by inhibiting raptor. Thus, we sought to investigate whether downregulation of the PI3K/Akt pathway influences viral replication by chemically inhibiting several pathway kinases. Huh-7 cells were preincubated with the PI3K inhibitor LY294002 (40 µM) or 300 nM of the novel clinically approved inhibitor copanlisib and subsequently infected with the YFV. Cell culture supernatants were collected 48 h after infection, and viral RNAs were quantified by RTqPCR (Fig. 8A). Inhibition of the PI3K by pre-incubation with LY294002 reduced viral replication by 2.44 log_10_ (p<0.0001) and by 1.9 log_10_ with copanlisib (p=0.0001). This indicated that the regulation of AMPK by the PI3K/Akt signalling cascade activity is crucial for viral replication (Fig. 8A). Similarly, treatment of Calu-3 cells with LY294002 inhibited SARS-CoV-2 replication by 2.63 log_10_ (p=0.002), supporting the importance of the PI3K activity (Fig. 8B).

**Figure 8.**
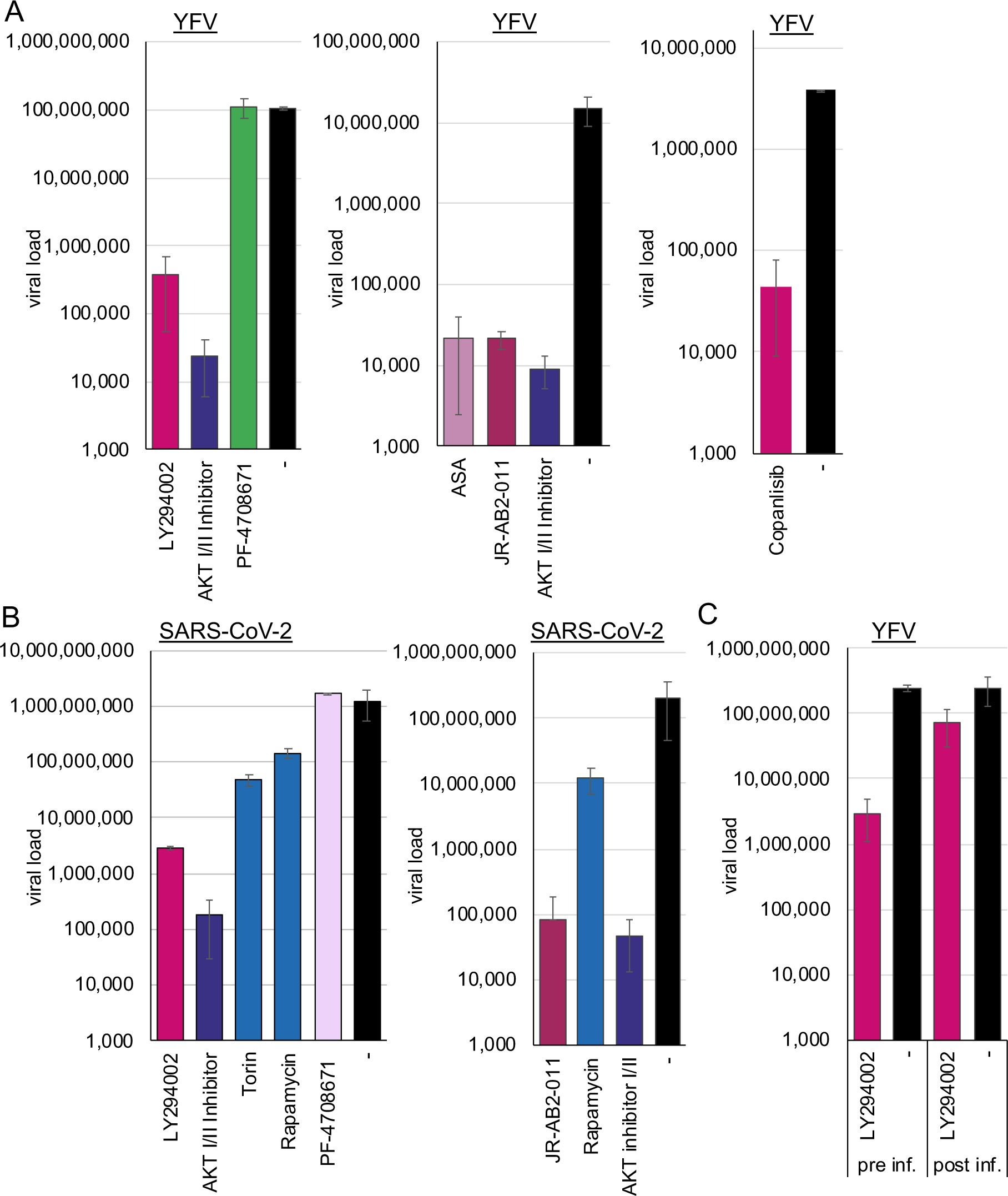
The PI3K, mTORC2, Akt, and mTORC1 are essential host factors for (A) flavivirus and (B) SARS-CoV-2 replication – (C) PI3K activity is required for YFV entry. **(A and C)** Huh-7 cells or **(B)** Calu-3 cells were incubated with inhibitors of PI3K (LY294002, copanlisib), Akt (AKT I/II), TORC1/2 (torin), TORC1 (rapamycin), S6 kinase (PF-4708671), and TORC2 (JR-AB2-011) and infected with **(A and C)** YFV or **(B)** SARS-CoV-2. Viral replication was analysed 72 h after infection**. (C)** YFV entry assays.

The PI3K is upstream of the Akt kinase, which controls the activation of NF-κB and, in turn, the expression of TXNIP. Thus, we sought to investigate whether the activity of Akt influences SARS-CoV-2 and flavivirus replication. The Akt kinase leads to the phosphorylation of mTOR. The Akt-I/II inhibitor blocks Akt kinase activity and, thus, autophosphorylation. Huh-7 and Calu-3 cells were incubated with 58 µM of Akt-I/II inhibitor and infected with YFV or SARS-CoV-2. Viral replication was determined at 48 h (YFV) or 72 h (SARS-CoV-2) after infection by RTqPCR (Fig. 8A and B). The Akt-I/II inhibitor treatment downregulated viral replication by more than 3 log_10_ log_10_ (YFV: 3.6 log_10_, p<0.00001; SARS-CoV-2: 3.84 log_10_, p=0.002). This result indicates that the activity of the Akt kinase is a crucial host factor for viral replication by inhibiting AMPK activation.

If AMPK activation and PI3K/Akt inhibition are two sides of the same coin, suppression of PI3K or Akt activity should also inhibit viral entry. Thus, we performed an entry assay similar to the experiments above with a PI3K inhibitor. Huh-7 cells were preincubated with LY294002 inhibitor for 2 h and infected or infected and treated with the inhibitor 2 h after infection. Viral RNAs were isolated after 48 h and quantified by RTqPCR (Fig. 8C). Inhibition of the PI3K before infection resulted in the suppression of flavivirus replication by 1.92 log_10_ (p<0.0001), indicating that PI3K activity and the PI3K/Akt pathway are essential for flavivirus replication. However, treatment after infection was ineffective (0.5 log_10_, p=0.009), suggesting that PI3K inhibition suppressed viral entry via increased AMPK activity. The importance of the Akt kinase activity was confirmed with TBEV and chikungunya entry assay (Fig. 2D and 3), where preincubation with the AKT I/II inhibitor suppressed viral replication by 3.3 log_10_ (TBEV, p=0.02) and 4.2 log_10_ (chikungunya, p=1.3×10^-5^).

Next, we sought to analyse the downstream effectors of Akt. The mTOR kinase’s activity controls major cellular pathways, such as NF-κB activation ^9^, cell metabolism, transcription, protein synthesis, autophagy and proteasome activity balance. Furthermore, mTORC1 regulates the activity of the lysosomal ATPase, which controls the lysosomal pH ^43^. The mTOR protein can be found in two distinct complexes, mTORC1 and mTORC2. The Akt kinase controls the first, while the second controls the Akt kinase independently of PI3K activity. First, we investigated the effects of inhibition of the mTOR activity by torin-1 and temsirolimus, a novel clinical-approved mTOR inhibitor. NCE cells were either preincubated for 2 h or TBEV infected and treated after 2 h with 10 µM temsirolimus, while Calu-3 cells were treated with 100 nM torin-1 and subsequently infected with SARS-CoV-2. Viral replication was determined with RTqPCR at 48 h (TBEV) or 72 h (SARS-CoV-2) (Fig. 2D and 8B) postinfection. The inhibition of both mTORC1 and −2 reduced SARS-CoV-2 replication (1.4 log_10_; p=0.003), showing that mTOR activity is required for viral replication (Fig. 8), while the TBEV entry experiments confirmed the inhibition of viral entry (1.4 log_10_, p=0.026). Experiments with rapamycin were performed to characterise further the mTORC complex, which inhibits SARS-CoV-2. Rapamycin selectively targets mTORC1. Incubation of Calu-3 cells with rapamycin inhibited viral replication (SARS-CoV-2: 0.9 log_10_; p=0.005), providing evidence that mTORC-1 activity is essential for viral replication (Fig. 8B). The connection of mTORC-1 to flaviviruses and SARS-CoV-2 replication can be explained by control of AMPK activity and, additionally, in the case of SARS-CoV-2, the mTOR-mediated acidification of the lysosomes.

Next, we analysed whether mTORC2 is required for SARS-CoV-2 and YFV replication since mTORC2 activates Akt independently of PI3K via Akt serine 473 phosphorylation. Calu-3 or Huh-7 cells were incubated with the specific mTORC2 inhibitor JR-AB2-011 (Fig. 8A and B). The cells were incubated with 250 µM JR-AB2-011 inhibitor and infected with YFV or SARS-CoV-2. Viral genome copies in the supernatants were quantified 48 h and 72 h after infection by RTqPCR. JR-AB2-011 suppressed YFV and SARS-CoV-2 replication by approximately 3 log_10_ (YFV: 2.84 log_10_, p=0.0003; SARS-CoV-2: 3.21 log_10_, p=0.002), indicating that mTORC2 plays a significant role in YFV and SARS-CoV-2 replication, which can result in Akt activation. Moreover, the activation of AMPK by ASA downregulated phosphorylation of Akt at serine 473 in non or SARS-CoV-2 infected Calu-3 cells, which was mediated by the mTORC2 complex, providing evidence that AMPK does not only control mTORC1 by phosphorylation of raptor but mTORC2 as well. This leads to the inhibition of viral replication since we have shown that Akt is essential for SARS-CoV-2 replication.

mTORC1 regulates the translation of specific heat-shock-related mRNAs by controlling the phosphorylation of the ribosomal S6 protein and the translation of 5’TOP-RNAs. Thus, we investigated whether the PI3K signalling cascade inhibits viral replication by downregulating the translation of viral RNA. The S6-kinase inhibitor PF-4708671 specifically suppresses S6 protein phosphorylation. Calu-3 and Huh-7 cells were incubated with 10 µM PF-4708671 and infected with YFV and SARS-CoV-2. Viral RNAs were isolated 48 h and 72 h after infection and quantified by RTqPCR (Fig. 8A and B). PF-4708671 did not influence viral replication, showing that S6 phosphorylation is dispensable for flavivirus or SARS-CoV-2 replication (SARS-CoV-2: −0.13 log_10_, p=0.17). However, AMPK activation by ASA downregulated S6 phosphorylation in Calu-3 and Huh-7 cells, which is expected when ASA leads to the dephosphorylation of Akt. We have shown that the S6 phosphorylation is not essential for SARS-CoV-2 replication. Thus, the ASA-mediated inhibition of S6 activity should not be responsible for the antiviral effect of ASA.

In summary, we have shown that active PI3K/Akt and mTORC2 pathways and the repression of AMPK/ULK1 are crucial for flavivirus, alphavirus, enteroviruses, SARS-CoV-2, and rabies lyssavirus entry and replication. This inhibition of receptor-mediated endocytosis and viral entry was independent of the AMPK activation pathway since ASA/SA, metformin, and the AMPK activator act on different cellular targets. In addition, our results suggest that the regulation of viral endocytosis is ULK1-dependent. Furthermore, the observed downregulation of EEs and LEs supports the reported AMPK-dependent upregulation of Rab4, an endosome recycling factor ^13^. The observed downregulation of LAMP-1 indicates that inhibiting EEs and LEs leads to reduced cellular lysosomes. Our findings suggest that AMPK/ULK1 activation inhibits receptor-mediated endocytosis and changes the number or composition of lysosomes. This central mechanism is a prime target for developing broad-spectrum antivirals. Finally, inhibiting rabies entry might lead to modifying the current postexposure prophylaxis, especially in the absence of antiserum with inexpensive and globally available drugs, such as ASA or metformin.

## Methods

### Viruses, antibodies, plasmids and chemicals

YFV strain: STAMARIL 17D-204; DENV type 2 strain ^44^, the patient-derived SARS-CoV-2 and the Chikungunya isolate have been described before ^45, 46^. *Wild-type* Rabies lyssavirus (015V-03645) was obtained from the FLI (Greifswald, Germany), the recombinant vaccine strain SPBN GAS from IDT Biologika (Germany). The patient-derived rubella, enterovirus 71, TBEV isolates were obtained from the virus diagnostics Würzburg, Germany and used with permission. All antibodies were obtained from Sigma-Aldrich (Taufkirchen, Germany), Cell Signaling (Leiden, Netherlands), and Thermo Scientific (USA). All chemicals and inhibitors were purchased from Sigma-Merck (Darmstadt, Germany), MedChemExpress (USA), and Roth (Karlsruhe, Germany). Activators: AMPK (CAS 849727-81-7), ULK1 (CAS 2101517-69-3).

### Western blotting analysis

For Western blot analysis, cells were washed with ice-cold PBS buffer and lysed in RIPA buffer (150 mM NaCl; 1% NP40; 12 mM sodium-desoxycholate, 3.5 mM SDS; 50 mM Tris pH 7) with sonification. Then Laemmli buffer (2% SDS, 10% Glycerol, 60 mM Tris, 0.01% (w/v) bromophenol blue, 37.5 μM β-Mercaptoethanol) was added. Proteins were separated by SDS PAGE (8-12%) and transferred to a nitrocellulose membrane using a wet blotting system (Carl Roth, Germany) using Towbin buffer (0.025 M TRIS 0.192 M Glycine, 20 % Methanol) for 1 to 3 h. Membranes were blocked with 5 % non-fat milk in PBS buffer for 20 min and incubated with respective antibodies overnight.

### Cellular proliferation assays

Direct automatic cell counting determined the proliferation of cells with and without the compounds. Cells were seeded on optical plates (CellCarrier-96, PerkinElmer) and counted before the experiments. Then, the compounds were added in decreasing concentrations, and the cells were incubated for three days. The cell numbers per well were determined using the PerkinElmer Ensight reader. Only compound concentrations that did not significantly reduce the cell number per well were used for antiviral assays.

### Viral infection, RNA isolation, detection, and viral load determination

The cells were incubated with the compounds either 2 h before or 2 h after infection and infected with the respective viruses. The medium was exchanged to remove inactive viruses, influencing genome copy determination with the medium containing the compounds (except AMPK or ULK1 activators). All infection experiments were performed in triplicate assays and repeated at least twice in independent experiments. After 48 or 72 h, 200 µl of the medium was collected, and viral genomes were purified with the High Pure Viral Nucleic Acid kit (Roche, Mannheim, Germany). Genome quantification of DENV, SARS-CoV-2 and TBEV was performed with the respective LightMix assay (TIBMolBiol), and YFV was done as described before ^47^. The provided standard was used for genome copy-number quantification using the LightCycler 480 II or Roche Lightcycler96 Software (Roche, Mannheim, Germany).

### Generation of Neural Organoids and PCLS

Neural organoids were generated from human induced pluripotent stem cells (hiPSCs), as previously described in detail ^48^. PCLS were prepared and infected as described ^22, 30, 46^.

### FACS staining and analysis

After 2 h treatment with AMPK or ULK, cells were washed with FACS buffer, and 1×10^6^ cells were resuspended in 50 µl of PBS supplemented with the primary antibodies and the viability dye on ice for 30 minutes. After that, the cells were washed with FACS buffer and counterstained with the corresponding conjugated secondary antibodies for 30 minutes on ice. The primary antibodies used: rabbit monoclonal anti-CD155 (1:100 dilution, clone: 4B3, Sigma-Aldrich) and mouse monoclonal anti-LDL receptor (1:100 dilution, clone: 2H7.1, Sigma-Aldrich). For live/dead staining, Fixable Viability Dye eFluor™ 780 were used (1:1000 dilution, cat. number: 65-0865-14, eBioscience). The secondary antibodies used: goat anti-rabbit Alexa Fluor 647 (1:100 dilution, polyclonal, cat. number: A-21244, Invitrogen) and goat anti-mouse PE (1:100 dilution, polyclonal, cat. number: 550589, BD bioscience). Samples were measured by an Attune NxT Flow Cytometer and analysed with FlowJo 10.10.0.

### Institutional Review Board Statement

Human lung lobes were acquired from patients undergoing lobe resection for cancer at Hannover Medical School. The use of the tissue for research was approved by the ethics committee of the Hannover Medical School and complies with the Code of Ethics of the World Medical Association (number 2701–2015). All patients gave written informed consent to use explanted lung tissue for research and publish the results. No sample tissues were procured from prisoners.

## Author Contributions

Conceptualisation, J.B., S.S-S., M.M., and M.L.; methodology, J.B., S.S-S, M.L., M.M; formal analysis, J.B., S.S-S; investigation, V.D.; V.L.R.; H.O.; M.T., K.N., J.B, H. A.; writing—original draft preparation, J.B.; writing—review and editing, V.D.; J.B., K.S., M.M., P.W., S.S-S., M.L. and M.S.; supervision, J.B.; M.S.; K.S.; M.M; P.W., M.L. and M.P.; project administration, J.B.; funding acquisition, J.B.; M.S.; K.S.; P.W., and M.L. All authors have read and agreed to the published version of the manuscript.

## References

1. Jeon, S.M. Regulation and function of AMPK in physiology and diseases. Exp Mol Med 48, e245 (2016).

2. Din, F.V. et al. Aspirin inhibits mTOR signaling, activates AMP-activated protein kinase, and induces autophagy in colorectal cancer cells. Gastroenterology 142, 1504–1515 e1503 (2012).

3. Hawley, S.A. et al. The ancient drug salicylate directly activates AMP-activated protein kinase. Science 336, 918–922 (2012).

4. Das, N. & Nebioglu, S. Vitamin C aspirin interactions in laboratory animals. J Clin Pharm Ther 17, 343–346 (1992).

5. Alvarez, C.E. On the origins of arrestin and rhodopsin. BMC Evol Biol 8, 222 (2008).

6. Qualls-Histed, S.J., Nielsen, C.P. & MacGurn, J.A. Lysosomal trafficking of the glucose transporter GLUT1 requires sequential regulation by TXNIP and ubiquitin. iScience 26, 106150 (2023).

7. Wu, N. et al. AMPK-dependent degradation of TXNIP upon energy stress leads to enhanced glucose uptake via GLUT1. Mol Cell 49, 1167–1175 (2013).

8. Waldhart, A.N. et al. Phosphorylation of TXNIP by AKT Mediates Acute Influx of Glucose in Response to Insulin. Cell Rep 19, 2005–2013 (2017).

9. Dan, H.C. et al. Akt-dependent regulation of NF-kappaB is controlled by mTOR and Raptor in association with IKK. Genes Dev 22, 1490–1500 (2008).

10. Miao, J., Zhou, X., Ji, T. & Chen, G. NF-kappaB p65-dependent transcriptional regulation of histone deacetylase 2 contributes to the chronic constriction injury-induced neuropathic pain via the microRNA-183/TXNIP/NLRP3 axis. J Neuroinflammation 17, 225 (2020).

11. Langemeyer, L., Frohlich, F. & Ungermann, C. Rab GTPase Function in Endosome and Lysosome Biogenesis. Trends Cell Biol 28, 957–970 (2018).

12. Gorvel, J.P., Chavrier, P., Zerial, M. & Gruenberg, J. rab5 controls early endosome fusion in vitro. Cell 64, 915–925 (1991).

13. Lee, J.O. et al. Metformin induces Rab4 through AMPK and modulates GLUT4 translocation in skeletal muscle cells. J Cell Physiol 226, 974–981 (2011).

14. Balderhaar, H.J. et al. The CORVET complex promotes tethering and fusion of Rab5/Vps21-positive membranes. Proc Natl Acad Sci U S A 110, 3823–3828 (2013).

15. Shvarev, D. et al. Structure of the endosomal CORVET tethering complex. Nat Commun 15, 5227 (2024).

16. Haas, A.K., Fuchs, E., Kopajtich, R. & Barr, F.A. A GTPase-activating protein controls Rab5 function in endocytic trafficking. Nat Cell Biol 7, 887–893 (2005).

17. Vonderheit, A. & Helenius, A. Rab7 associates with early endosomes to mediate sorting and transport of Semliki forest virus to late endosomes. PLoS Biol 3, e233 (2005).

18. De Luca, M. et al. RILP regulates vacuolar ATPase through interaction with the V1G1 subunit. J Cell Sci 127, 2697–2708 (2014).

19. Moser, T.S., Schieffer, D. & Cherry, S. AMP-activated kinase restricts Rift Valley fever virus infection by inhibiting fatty acid synthesis. PLoS Pathog 8, e1002661 (2012).

20. Glatthaar-Saalmuller, B., Mair, K.H. & Saalmuller, A. Antiviral activity of aspirin against RNA viruses of the respiratory tract-an in vitro study. Influenza Other Respir Viruses 11, 85–92 (2017).

21. Mazur, I. et al. Acetylsalicylic acid (ASA) blocks influenza virus propagation via its NF-kappaB-inhibiting activity. Cell Microbiol 9, 1683–1694 (2007).

22. Geiger, N. et al. Acetylsalicylic Acid and Salicylic Acid Inhibit SARS-CoV-2 Replication in Precision-Cut Lung Slices. Vaccines (Basel) 10 (2022).

23. Liu, Q. et al. Effect of low-dose aspirin on mortality and viral duration of the hospitalized adults with COVID-19. Medicine (Baltimore) 100, e24544 (2021).

24. Hampson, K. et al. Estimating the global burden of endemic canine rabies. PLoS Negl Trop Dis 9, e0003709 (2015).

25. Kosaka, T. et al. Identification of molecular target of AMP-activated protein kinase activator by affinity purification and mass spectrometry. Anal Chem 77, 2050–2055 (2005).

26. Zhou, G. et al. Role of AMP-activated protein kinase in mechanism of metformin action. J Clin Invest 108, 1167–1174 (2001).

27. Stein, B.D. et al. Quantitative In Vivo Proteomics of Metformin Response in Liver Reveals AMPK-Dependent and -Independent Signaling Networks. Cell Rep 29, 3331–3348 e3337 (2019).

28. Ianevski, A., Giri, A.K. & Aittokallio, T. SynergyFinder 3.0: an interactive analysis and consensus interpretation of multi-drug synergies across multiple samples. Nucleic Acids Res 50, W739–W743 (2022).

29. Diesendorf, V. et al. Drug-induced phospholipidosis is not correlated with the inhibition of SARS-CoV-2 - inhibition of SARS-CoV-2 is cell line-specific. Front Cell Infect Microbiol 13, 1100028 (2023).

30. Geiger, N. et al. Cell Type-Specific Anti-Viral Effects of Novel SARS-CoV-2 Main Protease Inhibitors. Int J Mol Sci 24 (2023).

31. Hoffmann, M. et al. Chloroquine does not inhibit infection of human lung cells with SARS-CoV-2. Nature 585, 588–590 (2020).

32. Kobayashi, K. & Koike, S. Cellular receptors for enterovirus A71. J Biomed Sci 27, 23 (2020).

33. Mendelsohn, C.L., Wimmer, E. & Racaniello, V.R. Cellular receptor for poliovirus: molecular cloning, nucleotide sequence, and expression of a new member of the immunoglobulin superfamily. Cell 56, 855–865 (1989).

34. Zhang, R. et al. Mxra8 is a receptor for multiple arthritogenic alphaviruses. Nature 557, 570–574 (2018).

35. Cong, H., Jiang, Y. & Tien, P. Identification of the myelin oligodendrocyte glycoprotein as a cellular receptor for rubella virus. J Virol 85, 11038–11047 (2011).

36. Koch, J. et al. TMPRSS2 expression dictates the entry route used by SARS-CoV-2 to infect host cells. EMBO J 40, e107821 (2021).

37. Avota, E. et al. The Manifold Roles of Sphingolipids in Viral Infections. Front Physiol 12, 715527 (2021).

38. Wang, S. et al. Fasting blood glucose at admission is an independent predictor for 28-day mortality in patients with COVID-19 without previous diagnosis of diabetes: a multi-centre retrospective study. Diabetologia 63, 2102–2111 (2020).

39. Cai, Y. et al. Fasting blood glucose level is a predictor of mortality in patients with COVID-19 independent of diabetes history. Diabetes Res Clin Pract 169, 108437 (2020).

40. Nikolic, J. et al. Structural basis for the recognition of LDL-receptor family members by VSV glycoprotein. Nat Commun 9, 1029 (2018).

41. Zhai, X. et al. LDLR is used as a cell entry receptor by multiple alphaviruses. Nat Commun 15, 622 (2024).

42. Geiger, N. et al. The Acid Ceramidase Is a SARS-CoV-2 Host Factor. Cells 11 (2022).

43. Ratto, E. et al. Direct control of lysosomal catabolic activity by mTORC1 through regulation of V-ATPase assembly. Nat Commun 13, 4848 (2022).

44. Wu, H. et al. Novel dengue virus NS2B/NS3 protease inhibitors. Antimicrob Agents Chemother 59, 1100–1109 (2015).

45. Kowalzik, S. et al. Characterisation of a chikungunya virus from a German patient returning from Mauritius and development of a serological test. Med Microbiol Immunol 197, 381–386 (2008).

46. Zimniak, M. et al. The serotonin reuptake inhibitor Fluoxetine inhibits SARS-CoV-2 in human lung tissue. Sci Rep 11, 5890 (2021).

47. Fischer, C. et al. Lineage-Specific Real-Time RT-PCR for Yellow Fever Virus Outbreak Surveillance, Brazil. Emerg Infect Dis 23, 1867–1871 (2017).

48. Worsdorfer, P., Rockel, A., Alt, Y., Kern, A. & Ergun, S. Generation of Vascularized Neural Organoids by Co-culturing with Mesodermal Progenitor Cells. STAR Protoc 1, 100041 (2020).

