## Supplemental Figures S1, S2 for "AMPK/ULK1 activation downregulates TXNIP, Rab5, and Rab7 and inhibits endocytosis-mediated entry of human pathogenic viruses"

Figure S1: AMPK activation inhibits entry of vesicular stomatitis virus

Figure S2. FACS gating Strategies

**Figure S1: AMPK activation inhibits entry of vesicular stomatitis virus.** Vero cells were treated either with 3 mM ASA 2h before or 2 h after infection. Viral RNAs were quantified 16 h after infection with CyberGreen and RTqPCR.

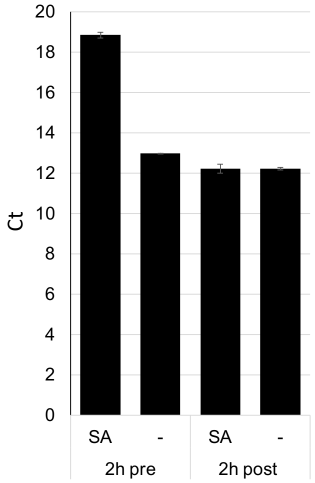


**Figure S2. FACS gating Strategies**

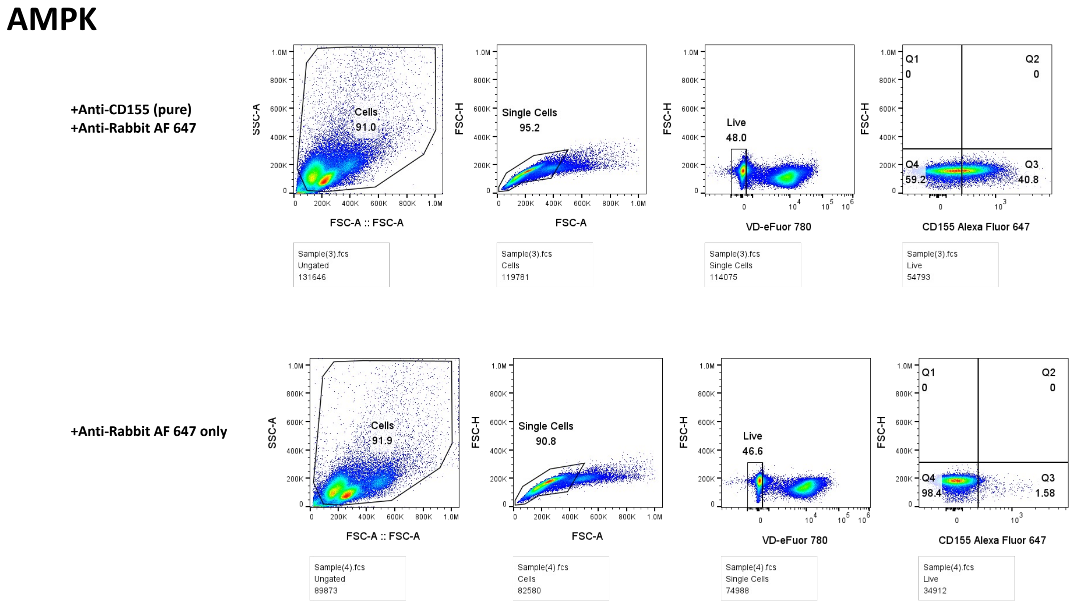

### Supplementary materials

#### ULK

**+Anti-CD155 (pure)**
**+Anti-Rabbit AF 647**

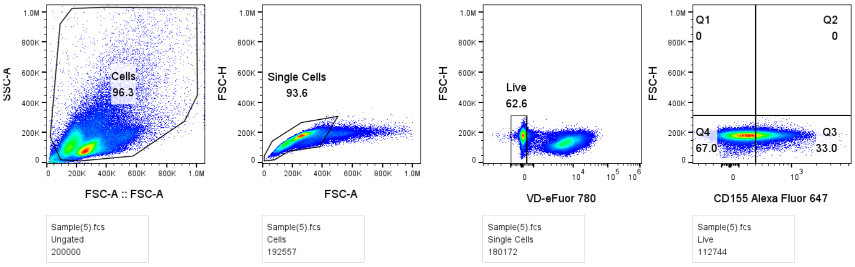


**+Anti-Rabbit AF 647 only**

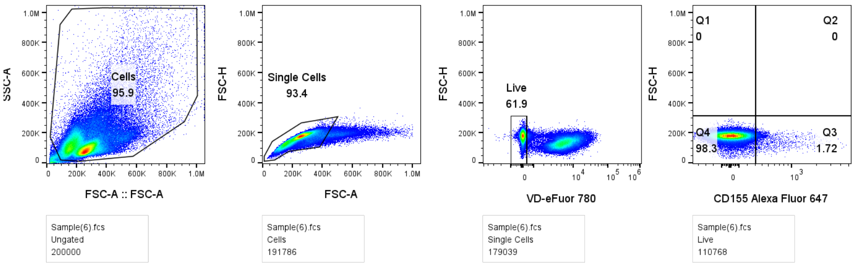


#### Untreated cells

**+Anti-CD155 (pure)**
**+Anti-Rabbit AF 647**

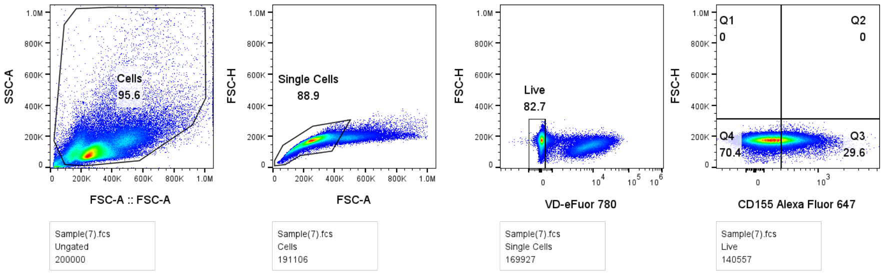


**+Anti-Rabbit AF 647 only**

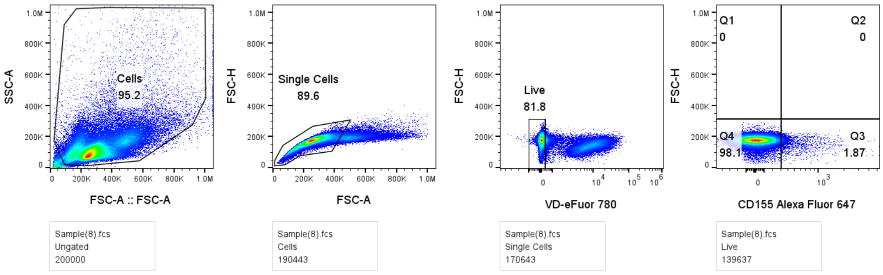

Supplementary materials

### Unstained cells

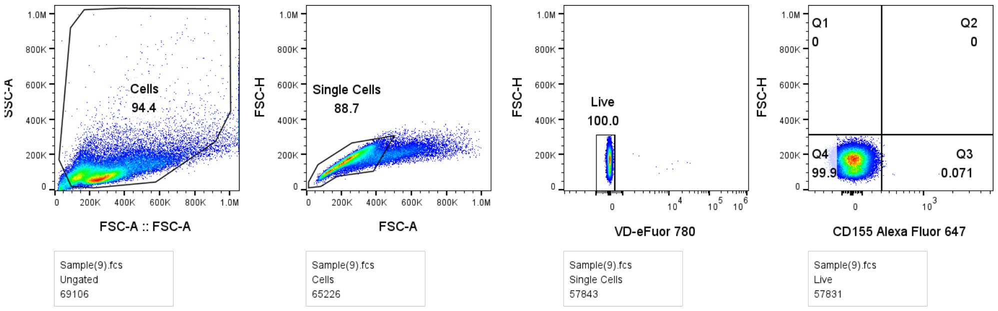


### Untreated cells

**+Anti-LDL (pure)**
**+Anti-Mouse PE**

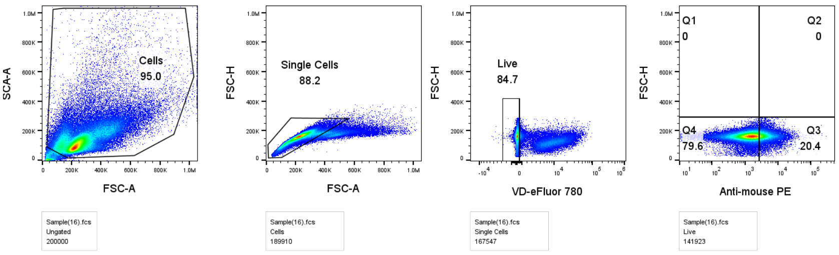


**+ Anti-Mouse PE only**

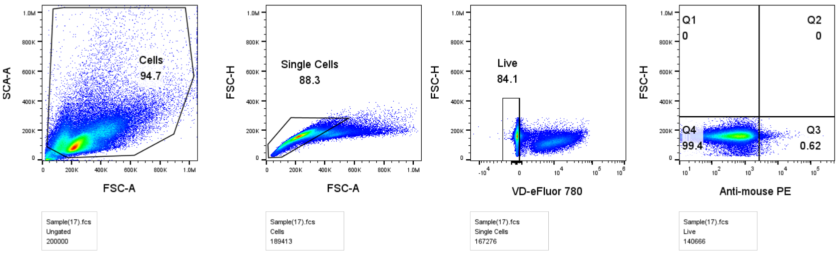

### Supplementary materials

#### Unstained cells

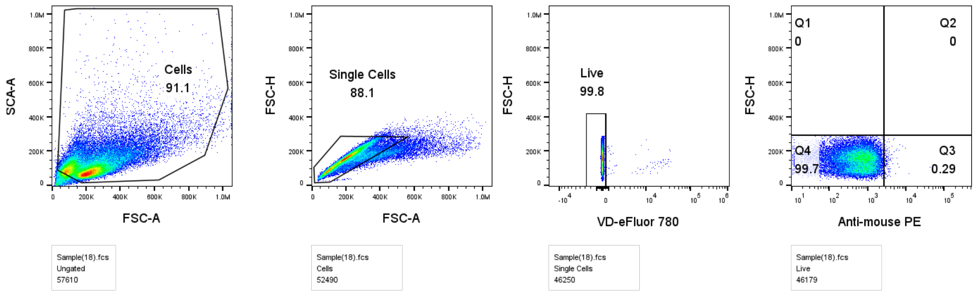


#### AMPK

**+Anti-LDL (pure)**
**+Anti-Mouse PE**

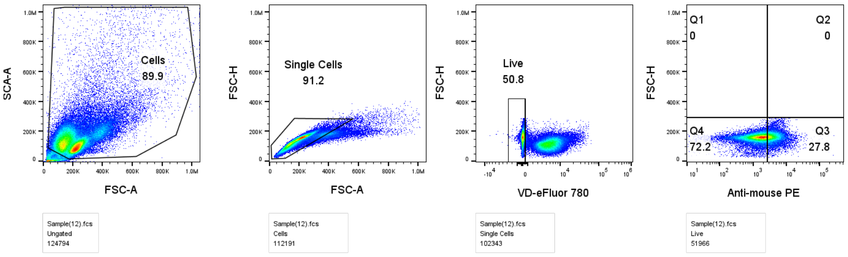


**+ Anti-Mouse PE only**

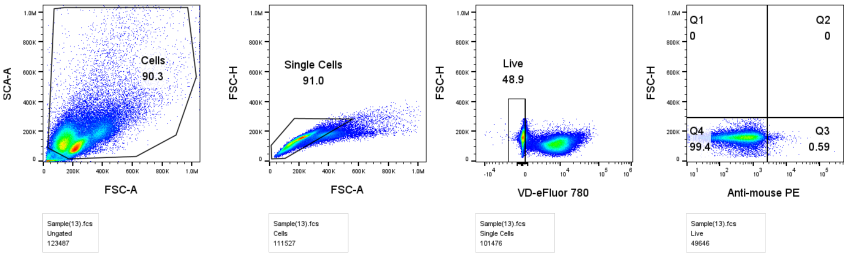

### Supplementary materials

#### ULK

**+Anti-LDL (pure)**
**+Anti-Mouse PE**

**+ Anti-Mouse PE only**

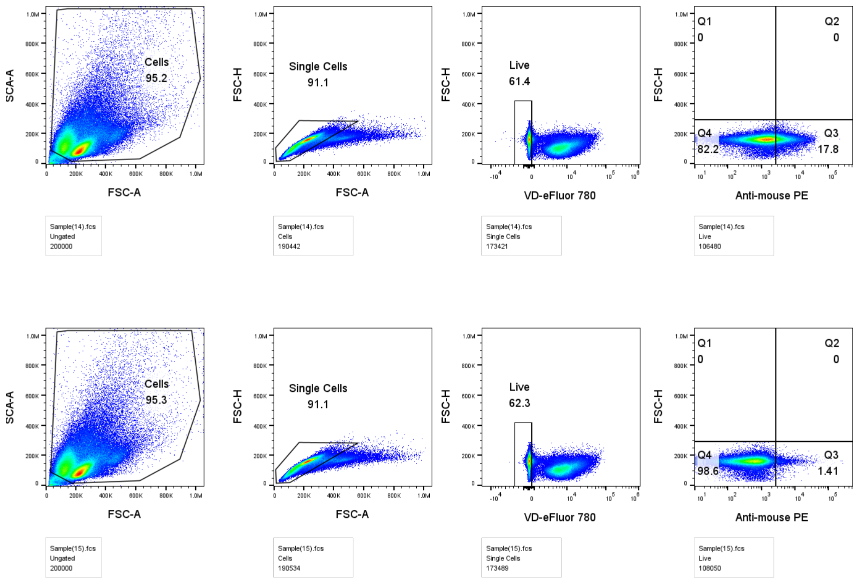
